# DYF-5 regulates intraflagellar transport by affecting train turnaround

**DOI:** 10.1101/2024.09.11.612404

**Authors:** Wouter Mul, Aniruddha Mitra, Bram Prevo, Erwin J.G. Peterman

## Abstract

Intraflagellar transport (IFT) coordinates the transport of cargo in cilia and is essential for ciliary function. CILK1 has been identified as a key regulator of IFT. The mechanism by which it acts has, however, remained unclear. In this study, we use fluorescence imaging and single-molecule tracking in the phasmid cilia of live *C. elegans* to study the effect of the CILK1 homolog DYF-5 on the dynamics of the IFT. We show that in the absence of DYF-5, IFT components accumulate at the ciliary tip. kinesin-II is no longer restricted to the proximal segment of the cilium but is present all throughout the cilium, while its velocity is different from that of OSM-3. The frequency of IFT trains is reduced and in particular retrograde trains were rarely observed. In the absence of DYF-5, retrograde transport is vastly reduced, resulting in the accumulation of IFT components at the tip and depletion at the base. The latter results in impeded anterograde train assembly, resulting in fewer trains with irregular composition. Our results show that DYF-5 plays a key role in regulating the turnarounds of IFT trains at the ciliary tip.

## Introduction

Many eukaryotic cells feature a structure called the cilium, which protrudes from the plasma membrane like an antenna. The primary cilium is an organelle that has a role in intracellular signalling, detecting extracellular cues including chemicals and light^1^. Cilia exhibit a wide range of functions and associated structures. All cilia are composed of an axoneme, a cylinder consisting of microtubules with their plus ends pointing outwards, away from the cell body. The axoneme is surrounded by the ciliary membrane, which is distinct from the plasma membrane both in phospholipid and membrane-protein composition^2,3^. The ciliary membrane is particularly enriched in transmembrane receptors.

The cilium relies on the intraflagellar transport (IFT) to preserve its structural and functional integrity. IFT is a bidirectional intracellular transport system, that moves cargo from the ciliary base to the tip (anterograde) and back from tip to base (retrograde). Anterograde movement is driven by kinesin-2 motors, retrograde transport by cytoplasmic dynein-2 (also known as IFT dynein)^4–7^. Cargo molecules (such as tubulin, transmembrane receptors or dynein) bind to so-called IFT trains, which consist of multiple copies of IFT-A and IFT-B protein sub-complexes, which are themselves composed of multiple proteins^8,9^. In the nematode *C. elegans,* IFT trains are driven in the anterograde of the cilium, for example at the end of dendrites in the case of *C. elegans* chemosensory cilia. Despite these challenges, IFT is continuous, with trains departing from the ciliary base every second or so, moving in one go towards the tip. At the tip, anterograde trains disassemble and reassemble into retrograde trains that move back towards the base in one continuous movement^10,11^. For IFT to work properly, strict regulatory mechanisms are required. Despite many studies, surprisingly little is understood about the regulation of IFT^12^. Several kinases have been described to be involved in the regulation of IFT. One of these is ciliogenesis associated kinase 1 (CILK1)^13,14^, which has been shown to be involved in regulating ciliary length and IFT-component localization. Knockouts of this protein in various model organisms have been shown to lead to reduced IFT velocity and abnormal accumulation of IFT components at the ciliary tip^13,15–17^. In some model organisms, these accumulations are shedded from the ciliary tip as vesicles^18–21^. How CILK1 regulates the IFT machinery is, however, not completely understood. One hypothesis is that it could regulate the interaction between motors and IFT train. This interpretation is based on observations that the velocities of both IFT trains and motors are altered in CILK1 knock outs^15,20,22,23^. Another suggestion is that it is involved in microtubule maintenance by regulation microtubule posttranslational modification and tubulin transport^15,24,25^.

Here we study the effect of CILK1 homolog DYF-5 on ciliary structure, IFT and motors in the phasmid chemosensory cilia of *C. elegans*, employing fluorescent labelling in combination with small-window illumination wide-field fluorescence microscopy (SWIM)^26^. Using this approach allows us to obtain single-molecule trajectories of IFT motors and IFT-train components in wild-type and *dyf-5* knock-out (KO) animals. We show that in the absence of DYF-5 both motors and IFT components accumulate at the tip of the cilium and that retrograde transport rate is dramatically reduced. We hypothesize that DYF-5 acts by regulating the turnaround of IFT train at the ciliary tip.

## Results

### Kinesin motor localisation in the presence and absence of DYF-5

In this study we investigate the regulation of IFT by the kinase DYF-5 in the phasmid chemosensory cilia of *C. elegans*. We visualized the distribution of kinesin motor proteins in *C. elegans* strains endogenously expressing fluorescent versions of kinesin-II (with an eGFP fused to the non-motor protein sub unit, KAP-1::eGFP) and OSM-3 (OSM-3::mCherry) in wild-type (WT) and *dyf-5* KO mutant backgrounds. We imaged the phasmid cilia in these animals using laser-illuminated epi-fluorescence microscopy. We time averaged fluorescence images and generated average fluorescence intensity profiles representing the distribution of motor proteins along a spline following the long axis of the cilium (Figure 1, A-D, first and third column). To provide insight into the heterogeneity between cilia and animals, we represent the distribution within multiple individual cilia as length-normalized heat maps (Figure 1, A-D, second column). These data show that in WT animals, kinesin-II is mostly localised in the proximal segment of the cilium peaking in concentration close the ciliary base (Figure 1A), while OSM-3 is present throughout the whole cilium, but at higher concentrations in the distal segments (Figure 1C). These localisation profiles are very consistent between animals (see standard deviation (Figure 1, A and C) and are similar to what has been published before^27,28^. In *dyf-5* animals, however, we observe rather different motor distributions. We note that in many of these mutant worms, ciliary structure appears abnormal (Figure 1, B-D, first column): ciliary length is highly variable and some cilia also show abnormal curvature, as has been observed before^15,17,24^. Focussing on the ciliary distribution of kinesin-II in *dyf-5* animals (Figure 1B), a striking observation is that kinesin-II is distributed throughout the whole cilium and no longer restricted to the proximal segment. Furthermore, in most cilia we observed kinesin-II accumulations at the ciliary tip (Figure 1B, heat maps). From the intensity distributions of individual cilia, we calculated the centre of mass (COM) and the variance around the centre of mass (Figure 1E). This analysis confirms the qualitative picture and shows that kinesin-II is consistently distributed differently in the absence of DYF-5.

**Figure 1.**
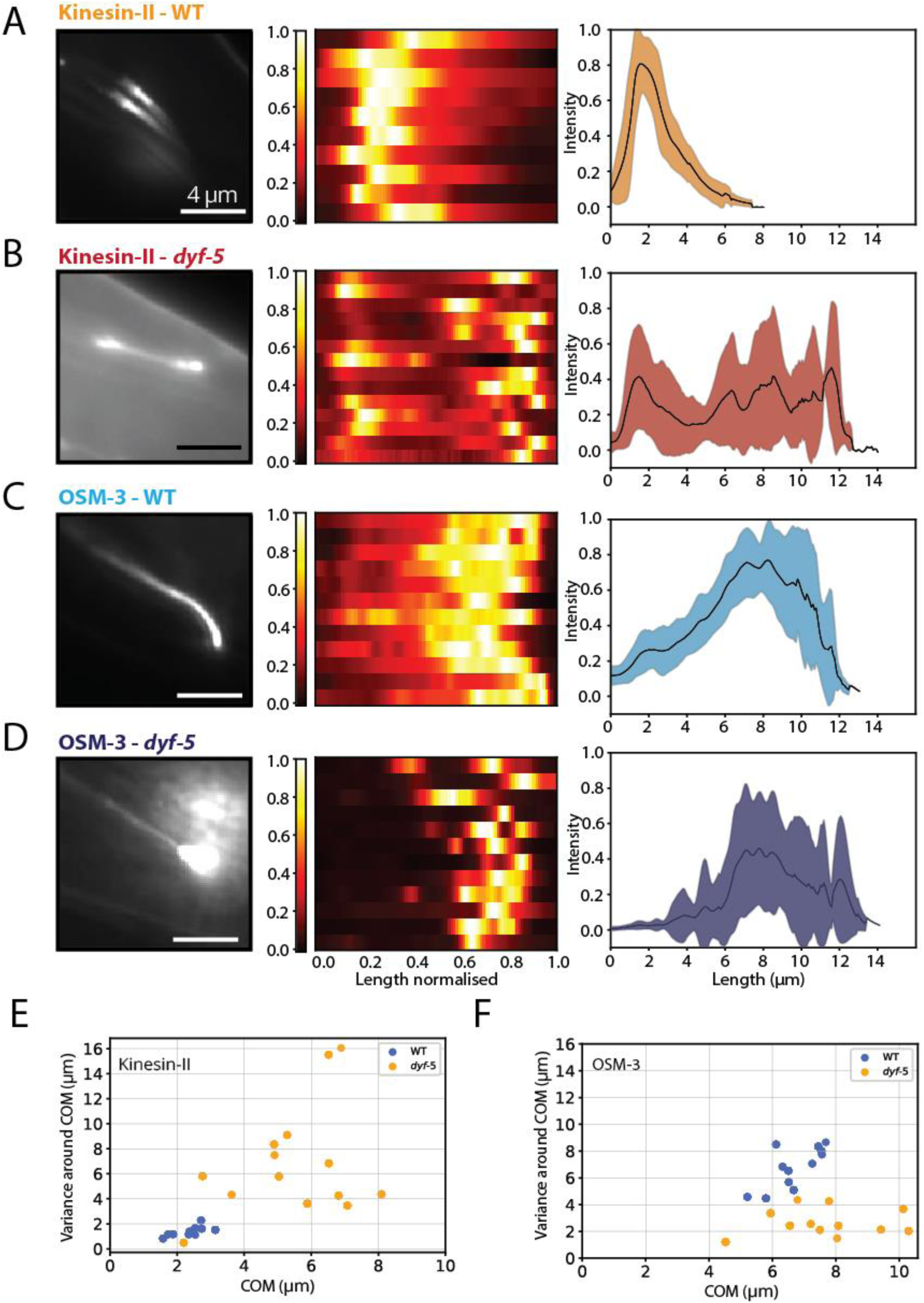
Impact of DYF-5 on cilia structure and kinesin-2 motor localization (A-D) Left: representative time-averaged fluorescence images; scale bar: 4 µm. Middle: heat maps of multiple individual cilia, length and intensity normalized. Look-up table of intensities shown on the left. Right: time averaged intensity profiles averaged over multiple cilia (as shown in middle column). Average is represented as solid black line, standard deviation in colour. (A) kinesin-II in the wild-type background (WT): KAP-1::eGFP, 10 cilia from 9 worms; (B) kinesin-II in *dyf-5* mutants: KAP-1::eGFP, 14 cilia from 14 worms; (C) OSM-3 in WT: OSM-3::mCherry, 12 cilia from 7 worms; (D) OSM-3 in *dyf-5*: OSM-3::mCherry,12 cilia from 12 worms. (E-F) Plots of the Variation around the centre of Mass (COM) of the intensity profiles of individual cilia against the COM for kinesin-II (E) and OSM-3 (F). The blue dots represent WT worms while the orange dots represent *dyf-5* KO worms.

We next focussed on the distribution of the other kinesin motor active in the phasmid cilia, OSM-3. At first sight, the difference in OSM-3 distributions in *dyf-5* and WT animals is less striking (Figure, 1C-D). We do, however, observe strong accumulations of OSM-3 at the ciliary tip, similar to kinesin-II (Figure 1D). These OSM-3 accumulations appear particularly intense and confined. The COM and variance around the COM of individual distributions clearly confirms the very intense and narrow OSM-3 accumulations at the ciliary tip, in almost all *dyf-5* animals (Figure 1F). Taken together, time-averaged bulk imaging of the ciliary distribution of the kinesin motors reveals three key effects of the absence of DYF-5 on the phasmid cilia: (i) cilia structure is disorganised and are often curved, (ii) kinesin-II is no longer restricted to the proximal segment, but distributed all along the cilium and (iii) both kinesin-II and OSM-3 accumulate in large numbers, locally and specifically at the ciliary tip.

### Bulk motor velocities and train frequencies in the presence and absence of DYF-5

We next focused on the dynamics of the kinesin motors in bulk imaging using the epifluorescence microscopy approach we developed before^28^. Typical kymographs of kinesin-II and OSM-3 in wild-type and *dyf-5* phasmid cilia are shown in figure 2. In both *dyf-5* kymographs (Figure 2C and D) static accumulations of the motors close to the ciliary tip can be observed, in agreement with the images and intensity profiles (Figure 1). We note that these accumulations are so intense that they precluded automated, quantitative analysis of velocities and intensities, as we have performed before on wild-type animals^28,29^. The kymographs do, however, allow a qualitative interpretation of the effect of the absence of DYF-5 on kinesin dynamics. Comparing the kinesins-II kymograph of *dyf-5* animals (Figure 2C) to wild type (Figure 2A) reveals that (i) kinesin-II velocity is substantially reduced (as evident from the less steep slope of the kymograph tracks); (ii) anterograde-moving kinesin-II accumulates in a static pool at the ciliary tip; (iii) the frequency of anterograde kymograph tracks is reduced, while retrograde tracks can hardly be discerned; (iv) the tracks are noticeable more blurry, which could indicate that kinesin-II motors repeatedly unbind from and rebind to IFT trains. For OSM-3 (Figure 2D), accumulations at the tip appear even more pronounced, as well as the decrease in anterograde track frequency and the almost absence of retrograde tracks. OSM-3 velocity appears hardly affected, in contrast to kinesin-II. This latter observation is remarkable, since it appears from the kymographs (Figure 2, C and D) that kinesin-II and OSM-3 do not travel together, at the same velocity in *dyf-5* animals, like they do in the proximal segment in wild type (Figure 2, A and B and in our previous work^28^).

**Figure 2.**
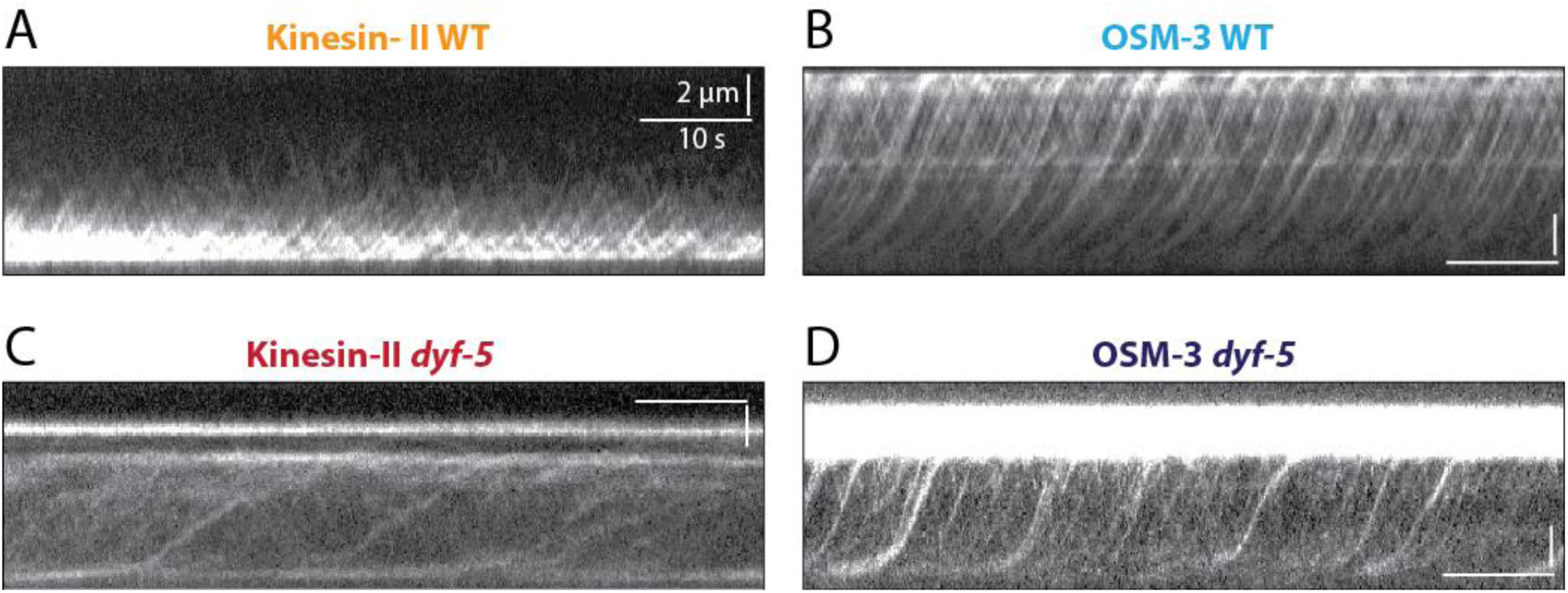
In the absence of DYF-5, bulk kinesin-II and OSM-3 velocities and frequencies are affected (A-D) Representative kymographs showing the bulk motility of kinesin-II and OSM-3 in WT and dyf-5 mutants; scale bars: 2 µm and 10 second. (A) Kinesin-II (KAP-1::eGFP) in WT, showing distinct tracks from base to handover zone and back. (B) OSM-3 (OSM-3::mCherry) in WT, showing clear tracks from base to tip and back again. (C) Kinesin-II (KAP-1::eGFP) in *dyf-5*, showing tracks extending to the ciliary tip. Kinesin-II velocity appears severely reduced. Furthermore, the anterograde and retrograde track frequencies appear lower. The intense horizontal line in the kymograph close to the ciliary tip is caused by a static pool of kinesin-II. (D) OSM-3 (OSM-3::mCherry) in *dyf-5*, showing tracks extending from base to the tip, but at a reduced frequency and almost no retrograde trains can be discerned. In addition, a very bright static pool is visible in the distal segment.

### Single-motor dynamics in the presence and absence of DYF-5

To shed more light on this observation that kinesin-II and OSM-3 velocities are differently affected by the absence of DYF-5, we performed single-molecule imaging of both motor proteins. To this end, we illuminated the cilium until almost all fluorescent proteins had photobleached. We then imaged single fluorescent kinesin proteins moving along the cilium using small-window illumination microscopy (SWIM), which allows relatively long term imaging at the single-molecule level, with an increased single-to-background ratio^26^. In many of the recorded movies, cilia were too curved to be in focus over their whole length within a single image plane and had to be discarded for further analysis. From the remaining image stacks, single-molecule kymographs were generated and single-molecule trajectories were extracted (Figure 3). Single-molecule tracks were filtered for anterograde movement and local point-to-point velocities were determined from all combined tracks of a given strain (Figure 3). Kinesin-II single-molecule kymographs from *dyf-5* cilia (Figure 3A and Supplementary Figure 1A-C) show a similar picture compared to wild type as bulk kymographs (Figure 2, A and C): (i) kinesin-II can be observed to move along the full length of the cilium; (ii) kinesin-II accumulates at the ciliary tip; and (iii) kinesin-II velocity is strongly reduced. Single-molecule velocity determination confirms that in wild type, kinesin-II gradually accelerates to ∼0.75 µm/s until it falls off IFT trains in the handover zone (2-4 µm into the cilium) (Figure 3A and Supplementary Figure 1A)^28^. This maximum velocity is higher than the intrinsic velocity of kinesin-II and is caused by the action of the faster OSM-3, which is involved in driving anterograde trains together with the slower kinesin-II in the handover zone^28^. Remarkably, in the absence of DYF-5, kinesin-II does not accelerate beyond ∼0.5 µm/s, which is believed to be close to its intrinsic velocity^28,30,31^. Similar velocities are observed throughout the cilium, beyond the transition zone.

**Figure 3.**
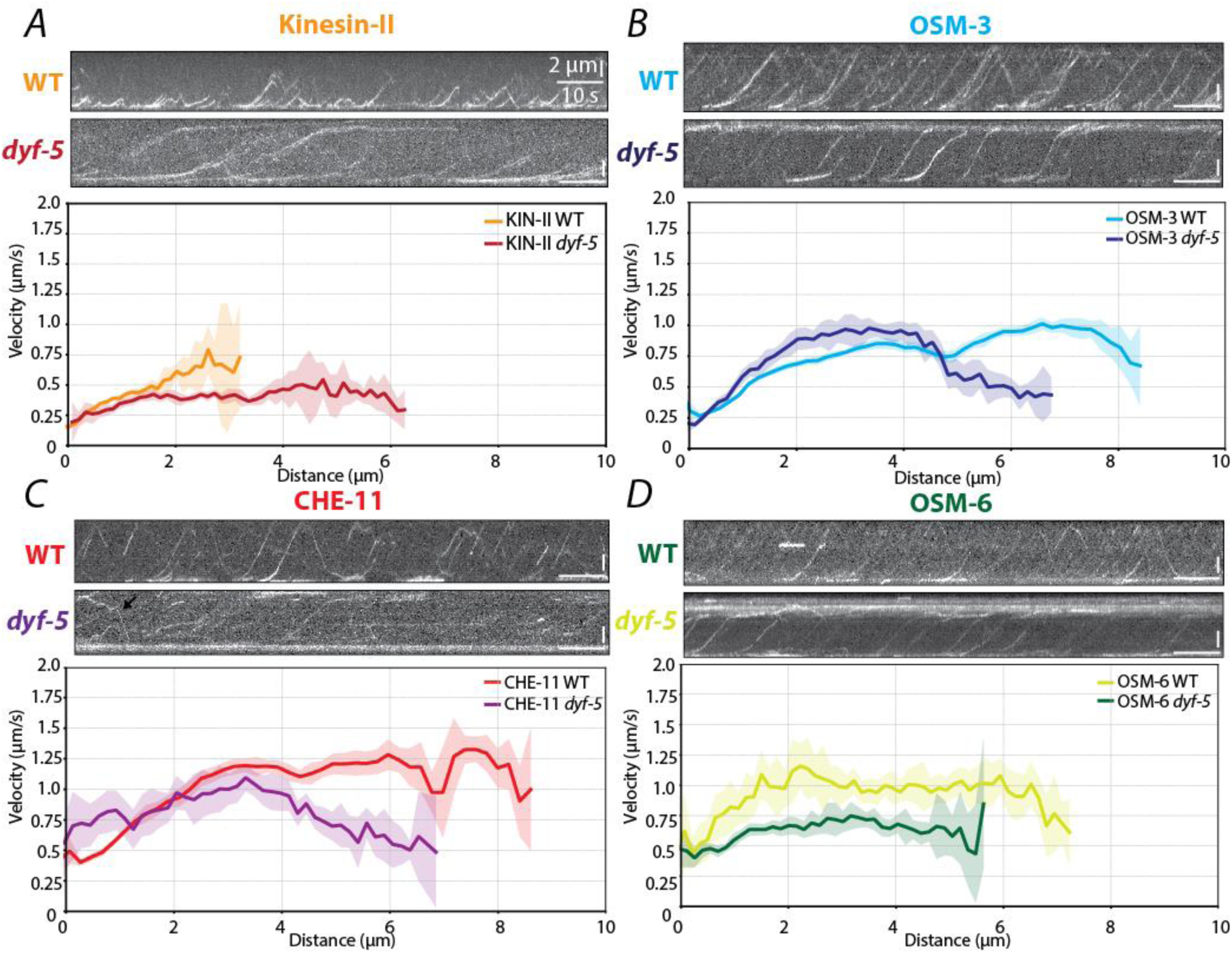
DYF-5 affects single-molecule velocities of IFT motors and IFT components (A-D) Representative single-molecule kymographs of WT (top), *dyf-5* KO (bottom) and binned average velocities along the length of the cilium (solid line ± SEM) for Kinesin-II (A), OSM-3 (B),CHE-11 (CHE-11::wrmScarlet part of the IFT-A complex) (C) and OSM-6 (OSM-6::eGFP part of the IFT-B) (D) (colours match the labels in the kymographs). Note that retrograde tracks are rarely visible in the *dyf-5* strains, but still happen occasionally (see e.g. the CHE-11 kymograph, black arrow). Scale bars: 2 µm and 10 seconds. (A) kinesin-II WT (top kymograph, orange; data from 15 cilia from 10 worms (9102 data point from 612 tracks)) and kinesin-II *dyf-5* (bottom kymograph, red; data from 10 cilia from 8 worms (5210 data point from 288 tracks)). (B) OSM-3 in WT (top kymograph, light blue; data from 19 cilia from 14 worms (27706 data point from 1083 tracks)) and OSM-3 in *dyf-5* (bottom kymograph, dark blue; data from 10 cilia from 9 worms (6713 data point from 276 tracks)). (C) CHE-11 WT (top kymograph, red; data from 10 cilia from 7 worms (9869 data point from 533 tracks)) and CHE-11 *dyf-5* (bottom kymograph, purple; data from 11 cilia from 9 worms (5860 data point from 356 tracks)). (D) OSM-6 WT (top kymograph, lime green; data from 9 cilia and 5 worms (4289 data point from 341 tracks)) and OSM-6 *dyf-5* (bottom kymograph, dark green; data from 10 cilia from 8 worms (5816 data point from 322 tracks))

**Figure 4.**
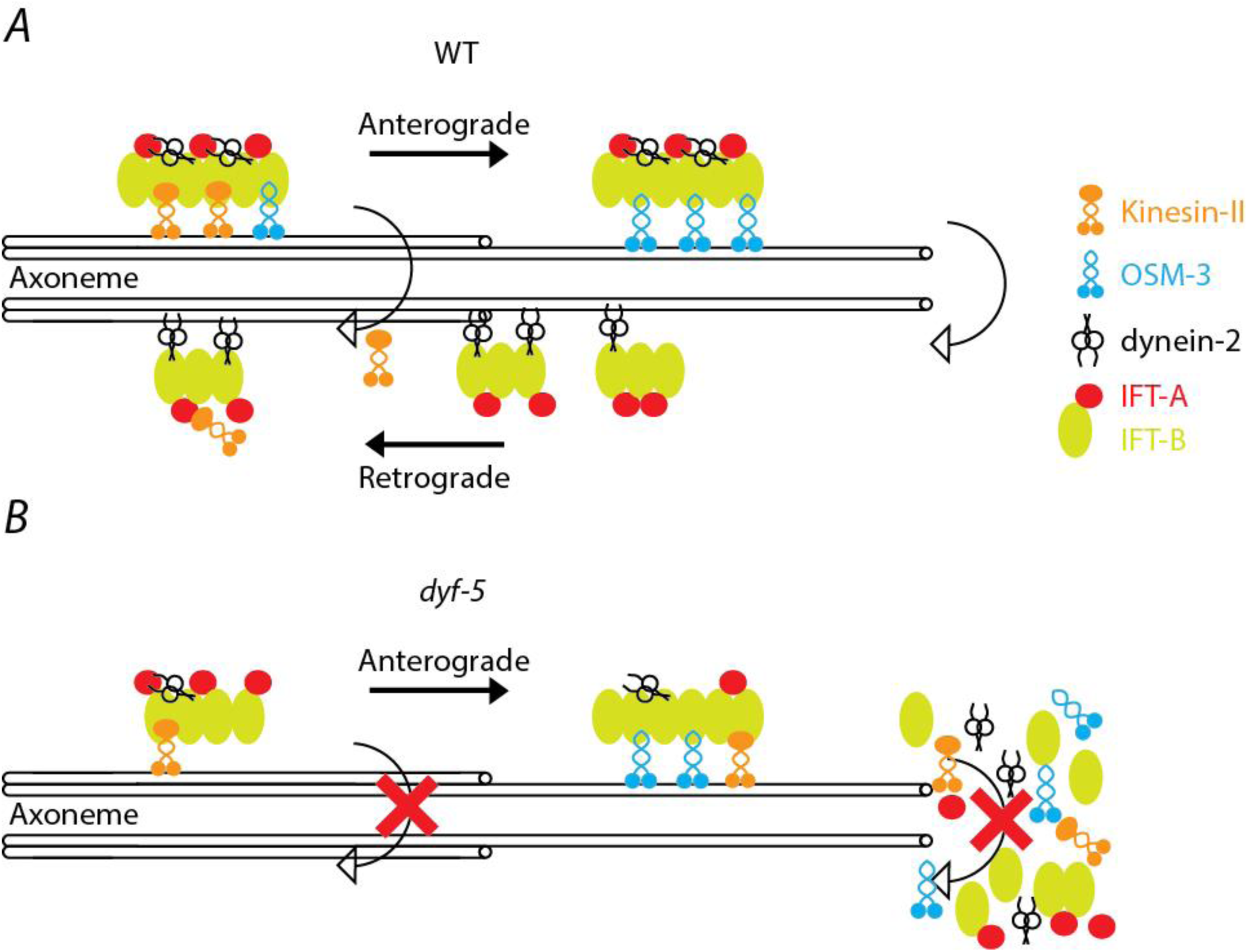
Illustration of IFT dynamics in the presence (WT) and absence of DYF-5 (*dyf-5)* (A) Schematic diagram illustrating IFT dynamics: IFT trains move in the anterograde direction from base to tip. At the tip, they reassemble into retrograde train, returning to the base. Indicated is a kinesin-II motor that switches from an anterograde to a retrograde train in the handover zone, where OSM-3 become the dominant anterograde motor on IFT trains taking care of the transport towards the ciliary tip. (B) In the absence of DYF-5, retrograde transport is inhibited, which has three consequences. (I) IFT components accumulate at the ciliary tip. (II) The frequency of retrograde trains is vastly reduced; therefore, kinesin-II has no retrograde trains to jump to. Instead, kinesin-II will rebind to another anterograde trains resulting in lower effective kinesin-II velocities, but also kinesin-II moving all the way towards the ciliary tip. (III) The reduced recycling of IFT components to the ciliary base results in a shortage of IFT components there, resulting in an increased variability of anterograde train composition, causing a wide distribution of motility characteristics.

OSM-3 single-molecule kymographs obtained in *dyf-5* cilia show a similar gradual acceleration in the ciliary proximal segment as in wild type (Figure 3B and Supplementary Figure 1B). This acceleration is caused by a gradual change in kinesin motor composition of trains, from mostly slower kinesin-II at the base and transition zone to relatively more faster OSM-3 further along the cilium, which has been described in detail before^28^. Kymographs suggest that OSM-3 velocity gradually decreases in the distal segment and static OSM-3 accumulations are clearly visible at the ciliary tip. Quantification of OSM-3 velocities (Figure 3B and Supplementary Figure 1D) confirms the qualitative picture: (i) a similar, gradual acceleration is observed from base to proximal segment, albeit that the observed maximum velocity is slightly lower in wild type (0.9 µm/s) than for *dyf-5* (1.0 µm/s) (ii) in *dyf-5*, but not in wild type, OSM-3 velocity drops in the distal segment to under 0.5 µm/s.

Focussing on the local velocities of both kinesins in wild type, we observe that both motors show a similar acceleration, to similar velocities, as observed before and as expected for two components traveling together on IFT trains. Remarkably, in *dyf-5* cilia, local OSM-3 and kinesin-II velocities appear uncoupled, reaching ∼1 µm/s and ∼0.5 µm/s, respectively between 2 and 4 µm along the cilium, which appears to match our bulk observations (Figure 2). We note that the variation in velocities of OSM-3 in particular is large in *dyf-5* animals (see for kymographs in Figure 3B and quantification of Figure 3B and Supplementary Figure 1D). Furthermore, a fraction of OSM-3 motors appears to travel at a different, lower velocity (Supplementary Figure 1D and Supplementary Figure 2 and 3).

### Single-molecule IFT-train dynamics in the presence and absence of DYF-5

This difference in velocities of the two kinesin motor proteins in *dyf-5* animals prompted us to investigate the integrity of IFT trains in this mutant strain. To this end we generated a nematode strain with both IFT-A (CHE-11::wrmScarlet) (Figure 3C) and IFT-B (OSM-6::eGFP) (Figure 3D) fluorescently labelled in the wild-type or the *dyf-5* mutant background. Kymographs and velocity profiles of WT animals (Figure3, C-D and Supplementary Figure 1, E-F) display that: both IFT-A and IFT-B move in one go from base to ciliary tip, gradually accelerating from the base and reaching a maximum velocity of 1.25 µm/s after about 3 µm. At the ciliary tip, the IFT-train components pause briefly before being transported back to the base as part of retrograde IFT trains. Kymographs taken in *dyf-5* animals (Figure3, C-D) show substantial, local accumulations of both IFT-A and IFT-B at the ciliary tip. Furthermore, the frequency of anterograde trains appears reduced. Retrograde tracks are even more rare but are not completely absent (see e.g. CHE-11 kymograph in Figure3C). Analysis of single-molecule velocities reveals that both IFT components move slightly slower in *dyf-5* animals (Figure 3, C-D and Supplementary Figure 1, E-H). The variation in velocities appears, however, substantially larger in *dyf-5* animals compared to wild type (Supplementary Figure 1, E-H): for example, a fraction of OSM-3 and CHE-11 appears to move at a higher velocity than the rest in *dyf-5* animals (Figure 3, C-D, bottom kymograph and Supplementary Figures 2 and 3). These results show that IFT trains have a more heterogeneous composition in the absence of DYF-5.

## Discussion

We have used fluorescence microscopy at the bulk and single-molecule levels to show that the absence of the CILK1 kinase homolog DYF-5 results in structural changes of the phasmid chemosensory cilia in *C. elegans*: cilia are more curved and the structure is disorganised. In addition, IFT is even more strongly affected. Kinesin-II is present along the whole length of the cilium and not only in the proximal segment and strong, static accumulations of kinesin-II, OSM-3, IFT-A and IFT-B are observed at the tips of most cilia. Furthermore, hardly any retrograde transport was observed. We also observed that IFT-train velocities appear to vary much more within a single cilium than in wild type.

Obtaining quantitative insight into the effect of the *dyf-5* knock-out proved challenging, in particular because of the large variability in length, shape and appearance between different cilia. Furthermore, imaging and analysis was hampered by the large, static accumulations of IFT components resulting in very bright spots in the fluorescence images, and by the increased curvature of the cilia resulting in parts of the cilia being frequently out-of-focus in images of the mutant worms. This made it very difficult to perform the automated analysis of in particular bulk kymographs we have used before^29^. Quantitative analysis of single-molecule images was less affected, since the high-intensity excitation required to reach the single-molecule regime caused photo bleaching of the accumulations.

The most prominent effect of the *dyf-5* mutation on the distribution of IFT motors and train components are the static accumulations we have observed at the ciliary tip (Figures 1,2 & 3). Similar observations have been made before in both *C. elegans* and other model organisms^16,17,32^. In IFT-dynein or IFT-A mutants, similar tip accumulations have been seen^33–36^. Furthermore is has been reported that dynein accumulates at the ciliary tip^15^. In line with those observations, in our *dyf-5* data retrograde transport is severely reduced. We have observed before that hyperosmotic stimulation of the cilia results in abrupt inhibition of the formation of retrograde IFT trains at the ciliary tip, resulting in similar accumulations of IFT components at the tip^37^. In those experiments, retrograde transport restarts almost immediately after switching back to isotonic buffer conditions and the accumulations disappear within seconds. We have not seen clear differences in hyperosmotic response in *dyf-5* mutant worms compared to wild type (data not shown), indicating that there is likely some redundancy in regulating the response. Taken together this suggests that DYF-5 plays a key role in the turnaround of IFT trains from anterograde to retrograde at the ciliary tip, but it is at this point unclear whether this role is in the disassembly of anterograde trains, the reassembly of retrograde ones and/or the activation of IFT dynein. In human cells lacking DYF-5 homolog CILK1, IFT-component accumulations have been observed to be shed from the tip in vesicles^21,38,39^. We have not directly observed shedding of vesicles from ciliary tips in *dyf-5* worms, which might be a consequence of the ciliary tips being rarely in the same focal plane as the rest of the cilium, due to the increased curvature of cilia in this strain. Occasionally, however, we did observe puncta of fluorescence beyond the ciliary tip, which could be vesicles stuck in the phasmid channels. We observed that the velocities of the IFT components slows down to 0.5 µm/s in the distal segment, near the ciliary tip, the location of the static accumulations. It could well be that these observations are correlated and that the low velocity is an effect of the high density of accumulated proteins, slowing down the trains by a crowding effect (Supplementary Figure 2).

Another striking observation in *dyf-5* worms is that kinesin-II is not restricted to the proximal segment (as it is in wild type) but is distributed all along the cilium. We note that we have observed a similar relocation of kinesin-II upon hyperosmotic stimulation of the chemosensory cilia, which halts the departure of retrograde IFT trains^37^. We believe that this redistribution is a direct consequence of the lack of retrograde transport in *dyf-5* animals. In wild type, kinesin-II gradually falls off anterograde IFT train beyond the transition zone and is picked up, after a short period of free diffusion, in most case by retrograde trains and transported back to the base^28,40^. In *dyf-5* cilia, kinesin-II likely still falls off anterograde trains, but due to the absence of retrograde trains, it is not transported back to base. It can only rebind to another anterograde train and is transported as cargo or as driver of IFT further along the cilium. This hypothesis is supported by the observation that kinesin-II kymograph tracks appear blurrier in the absence of DYF-5. We have before observed kinesin-II rebinding from anterograde to anterograde trains in wild type^40^. It could very well be that in *dyf-5*, kinesin-II alternates between episodes of being bound to anterograde IFT trains and of being free in solution. This would result in episodes of directed, anterograde motion and diffusion respectively, as we have observed before in wild type^40,41^. In this picture, kinesin-II motion would only be partly coupled to that of the rest of the IFT trains, including OSM-3, which is in line with our observation that the velocity of kinesin-II does not exceed 0.5 µm/s and is slower than OSM-3 and the IFT-particle components.

For the other train components studied (OSM-3, IFT-A and IFT-B) we observed larger variability in velocities than in wild type. Furthermore, the velocities of kinesin-II, OSM-3, IFT-A and IFT-B differ in *dyf-5* worms. For example, the velocities of OSM-3 and OSM-6 are marginally off for the whole length of the cilium. These differences in velocity are in stark contrast to wild type (Figure 3^28^), where these train components move at nearly identical position-dependent speeds, which is a signature of a well-defined location-dependent train composition and all these components being almost always attached to IFT trains and only very briefly moving independently. We interpret these different (spreads in) velocities in *dyf-5* worms to indicate that the composition of trains is much more variable in these worms. This might be caused by a scarceness of IFT components at the base owing to the almost complete absence of retrograde transport. Depending on the exact composition of a given train, it might move faster, slower or more variable and even fall apart along the way (Supplementary figure 2). Effects on IFT-train velocity in *dyf-5* worms have been observed before and interpreted to indicate that OSM-3 moves independently of IFT trains in these mutant strains. In different knock-out lines, for example affecting the BBSome, different IFT-A and IFT-B motility has been interpreted to indicate that they move independently from each other in these strains^42^. We show here that that IFT components moving separately is most likely not the case in these mutant strains. Most likely, these mutations cause a large variability in train composition, which subsequently results in different motility characteristics of individual trains, depending on their specific composition.

In summary, we have investigated here the effect of the kinase DYF-5 –the *C. elegans* homolog of human CILK-1– on IFT dynamics. Our results indicate that DYF-5 is required for the directional switches of IFT trains from anterograde to retrograde at the ciliary tip. Absence of DYF-5 results in accumulation of IFT components at the tip and depletion of components at the base. Over time, this depletion at the base appears to result in newly formed IFT trains with less uniform composition and disturbed motility.

## Supplementary figures

**Supplementary figure 1.**
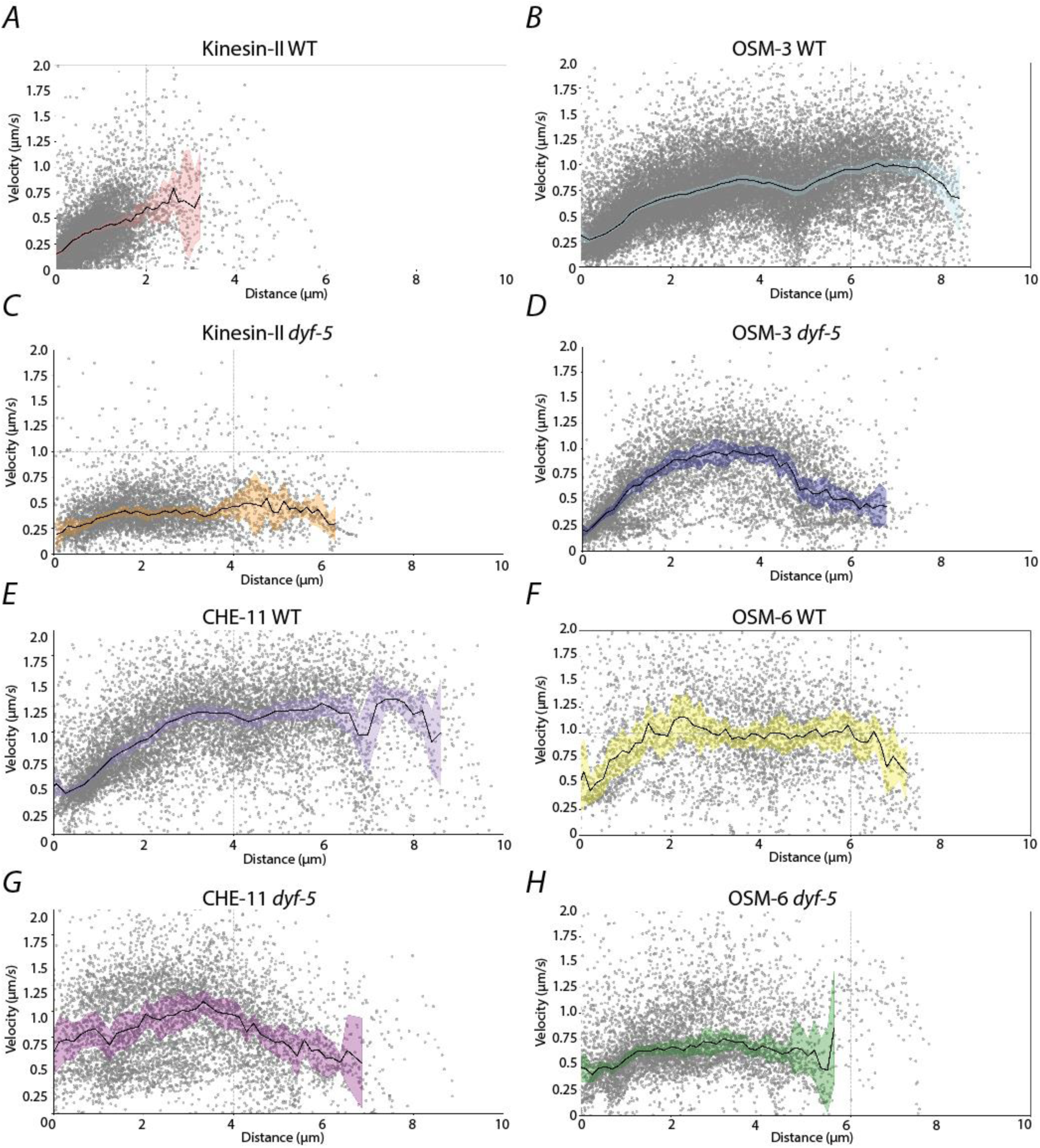
Underlying point-to-point velocity distribution along the ciliary length for different IFT components. (A-H) individual plots of the binned average velocities along the ciliary length (solid line ± SEM). Grey dots represent individual data points. Same data as in figure 3. (A) Kinesin-II in WT background (orange; data from 15 cilia from 10 worms (9102 data point from 612 tracks)). (B) OSM-3 in WT background (light blue; data from 19 cilia from 14 worms (27706 data point from 1083 tracks)). (C) kinesin-II in *dyf-5* KO background (red; data from 10 cilia from 8 worms (5210 data point from 288 tracks)). (D) OSM-3 in *dyf-5* KO background (dark blue; data from 10 cilia from 9 worms (6713 data point from 276 tracks)). (E) CHE-11 in WT background (red; data from 10 cilia from 7 worms (9869 data point from 533 tracks)). (F) OSM-6 in WT background (lime green; data from 11 cilia from 9 worms (5860 data point from 356 tracks)). (G) CHE-11 II in *dyf-5* KO background (purple; data from 9 cilia and 5 worms (4289 data point from 341 tracks)). (H) OSM-6 II in *dyf-5* KO background (dark green; data from 10 cilia from 8 worms (5816 data point from 322 tracks)).

**Supplementary figure 2.**
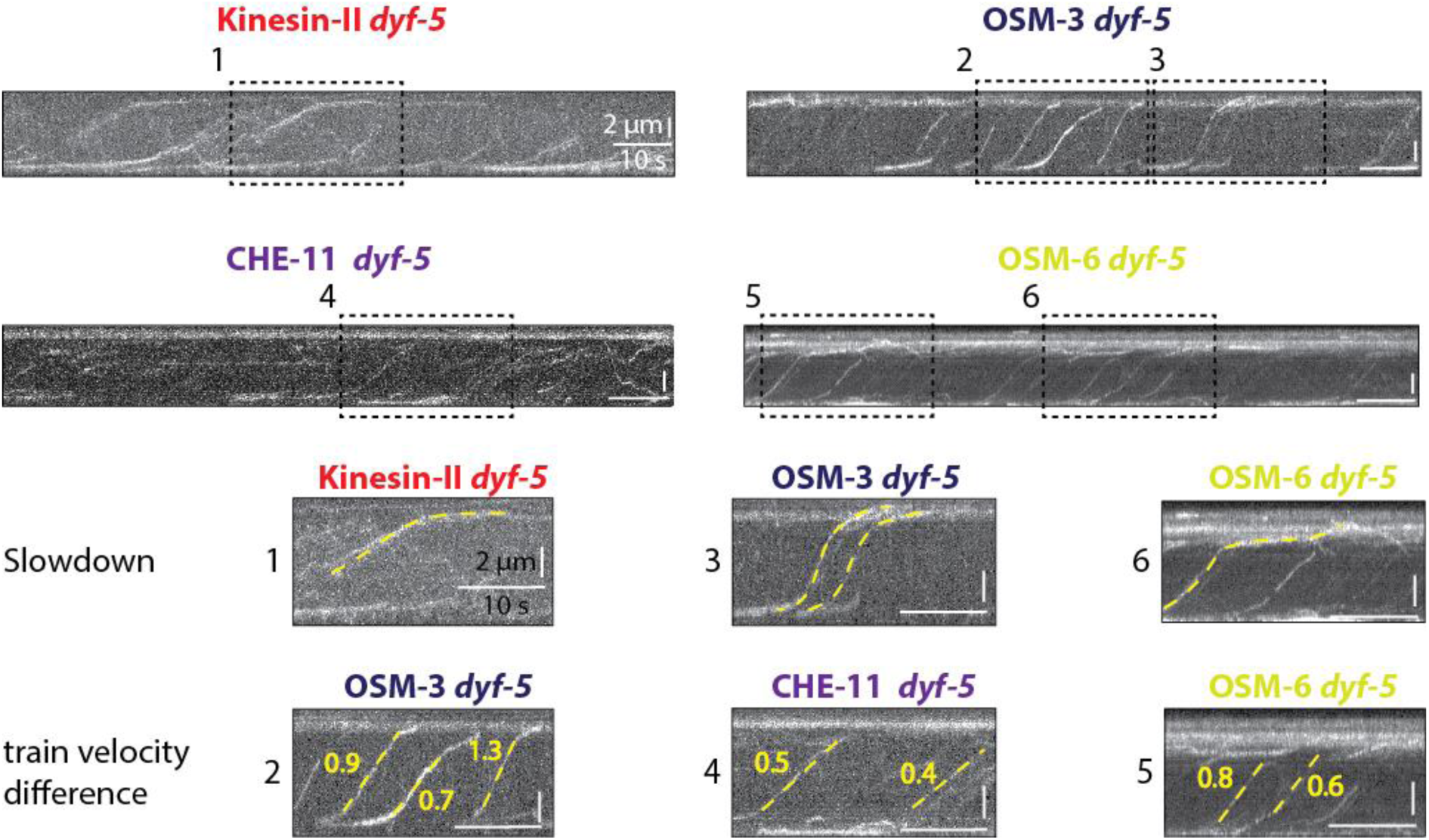
Exemplary events in kymographs obtained from single-molecule imaging of different IFT-components in dyf-5 mutants. Top, representative single-molecule kymographs of Kinesin-II, OSM-3, CHE-11 and OSM-6 in the *dyf-5* KO background. Zooms of the boxes indicated in the top kymographs, showing examples of slowdowns at the location of accumulations and examples of velocity changes of IFT train in the same cilium. Numbers indicate the average velocity of the tracks in µm/s. Scale bars: 10 seconds and 2 µm. Kymographs are the same as in figure 3.

**Supplementary figure 3.**
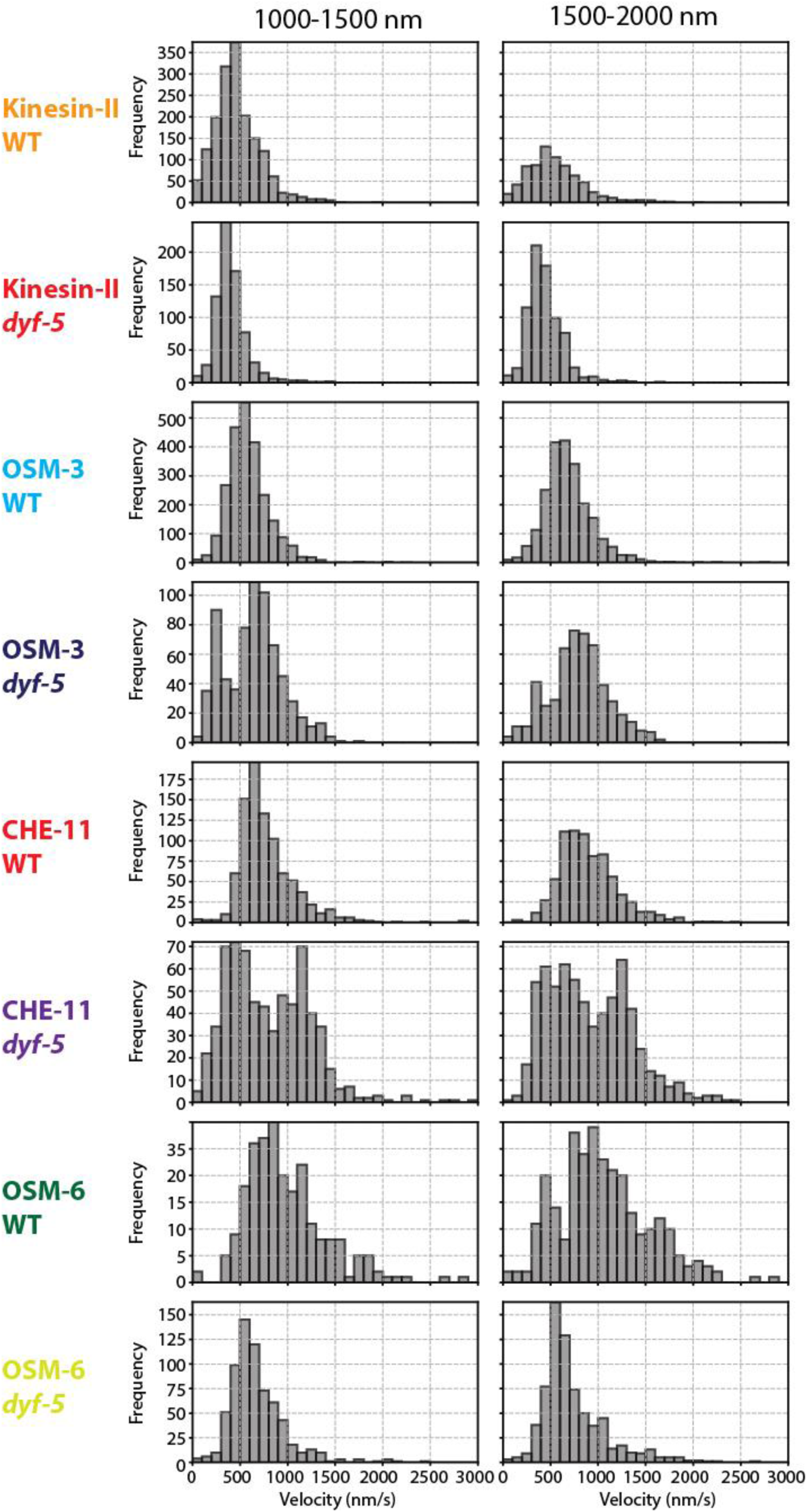
Distributions of point-to-point velocities obtained from single-molecule tracks of different IFT components. Histograms showing the distribution of single-molecule velocities of Kinesin-II, OSM-3, CHE-11, and OSM-6 in WT and *dyf-5* KO backgrounds. The distributions are displayed in two segments: (i) 1000-1500 nm from the base of the cilia (left); (ii) 1500-2000 nm from the base of the cilium. For Kinesin-II, the WT and *dyf-5* KO backgrounds show 1677 and 745 velocities respectively (1000-1500 nm), and 730 and 764 velocities (1500-2000 nm). For OSM-3, the WT and *dyf-5* KO backgrounds show 2415 and 2199 velocities respectively (1000-1500 nm), and 683 and 518 velocities (1500-2000 nm). For CHE-11, the WT and *dyf-5* KO backgrounds show 882 and 755 velocities respectively (1000-1500 nm), and 671 and 657 velocities (1500-2000 nm). For OSM-6, the WT and *dyf-5* KO backgrounds show 230 and 277 velocities respectively (1000-1500 nm), and 676 and 716 velocities (1500-2000 nm).

**Supplementary table 1.** *C. elegans* strains used in this study.

| Strain | Name used | Genotype | Source |
| --- | --- | --- | --- |
| EJP13 | Kinesin-II::eGFP | kap-1(ok676) III; vuaSi1 [pBP20; Pkap-1::kap-1::eGFP; cb-unc-119(+)] IV | Prevo *et al.*, 2015 |
| EJP28 | Kinesin-II::eGFP<br>$\Delta$ dyf-5 | dyf-5 (ok1177) I; kap-1(ok676) III; vuaSi1 [pBP20; Pkap-1::kap-1::eGFP; cb-unc-119(+)] IV | This study (MosSCI) |
| EJP16 | OSM-3::mCherry | vuaSi2 [pBP22; Posm-3::osm-3::mCherry; cb-unc-119(+)] II; unc-119(ed3) III; osm-3(p802) IV | Prevo *et al.*, 2015 |
| EJP43 | OSM-3::mCherry<br>$\Delta$ dyf-5 | dyf-5 (ok1177) I; vuaSi2 [pBP22; Posm-3::osm-3::mCherry; cb-unc-119(+)] II; unc-119(ed3) III; osm-3(p802) IV | This study (MoSCI) |
| Phx8141 | OSM-6::eGFP<br>CHE-11::wrmSCarlet | osm-6(syb7957[osm-6::eGFP]V,che-11(syb8141[che-11::wrmScarlet])V | Mitra *et al.*, 2024<br>(made by sunyBiotech) |
| PHX8631 | OSM-6::eGFP<br>CHE-11::wrmSCarlet<br>$\Delta$ dyf-5 | dyf-5 (ok1177) I; osm-6(syb7957[osm-6::eGFP]V,che-11(syb8141[che-11::wrmScarlet])V | This study(sunyBiotech) |

## Materials and methods

### *C. elegans* strains

*C. elegans* strains were cultured according to standard methods^43^, on NGM plates seeded with HB101 *E. coli*, at 20 ℃. Mutant worm lines were generated with Mos-1 mediated single-copy insertion (MosSCI)^44^ or with CRISPR/cas9^45^. The strains used in this study are listed in Supplementary table 1.

### Fluorescence imaging

Fluorescence imaging was performed as described before^46^ and for single-molecule imaging SWIM was used^26,41,47^. In short, young hermaphrodite adult *C. elegans* were sedated in 5 mM levamisole in M9 for a minimum of 10 minute. Then they were sandwiched between a coverslip and a 2 % agarose (M9) pad, sealed with VALAP^48^ followed by an hour of incubation before imaging. Images were acquired using a custom-built laser-illuminated wide-field fluorescence microscope, as described previously^46^. In short, optical imaging was performed using an inverted microscope body (Nikon Ti E) with a 100x oil immersion objective (Nikon, CFI Apo TIRF 100x, NA 1.49) in combination with an EMCDD camera (Andor, iXon 897) controlled using MicroManager software (v1.4). 491 nm and 561 nm DPSS lasers (Cobolt Calypso and Cobolt Jive, 50 mW) were used for laser illumination. Laser power was adjusted using an acousto-optic tuneable filter (AOTF, AA Optoelectronics). For SWIM, the beam diameter was changed using an iris diaphragm (Thorlabs, SM1D12, ø 0.8-12 mm), placed roughly at the back-focal plane of the objective. The beam width of the excitation beam was ∼30 μm (2σ of the Gaussian width). The aperture size of the diaphragm was adjusted manually to change the width of the beam, with a minimum beam diameter of ∼7 μm in the sample. Fluorescence light was separated from the excitation light using a dichroic mirror (ZT 405/488/561; Chroma) and emission filters (525/50 and 630/92 for collecting fluorescence excited by 491 nm and 561 nm, respectively; Chroma). Samples were typically imaged at 5.3x preamplifier gain and 300 EM gain with 10MHz ADC readout. The pixel size of acquired images was 80 nm × 80 nm (the field of view was cropped, imaging only the illuminated region). To image ensemble dynamics and obtain average fluorescence-intensity profiles, a cilia pair was imaged with a low intensity 491 nm or 561 nm beam (∼0.1 mW/mm^2^) for 5 minutes at 6.6 fps. To perform single-molecule imaging, the sample was illuminated with a high intensity (∼10 mW/mm^2^), photobleaching the signal in the cilia pair for roughly 10 min before starting acquisition. This approach allows us to completely photobleached the fluorescent signal from the accumulations at the distal part of the cilia and visualize only the dynamics of fresh (unbleached) molecules entering the illuminated region. The frame rate was 6.6 frames per second (150 ms exposure time).

### Intensity profiles

For the fluorescence intensity profiles, movies were acquired for 5 minutes with 150 ms exposure time, at low laser intensity (∼0.1 mW/mm^2^). Time-averaged projections were made by averaging the frames in the complete movie. Followed by this, a manual segmented line with a linewidth of 7pixels was drawn and the intensity along this line was determined using ImageJ. The start of the segmented line was determined by the user to be just before the ciliary base. The background intensity was determined next to the cilium and subtracted from the raw intensity of the cilium. Intensities were normalized to the maximum intensity. The average cilium-intensity profiles were obtained by averaging over multiple cilia, with error provided by the standard deviation (typically N = 10 cilia). For cilia in EJP28 (kinesin-II::eGFP in *dyf-5* mutants), since the cilia are highly curved (never completely in a single plane of focus) and the signal is relatively weak, we acquired Z-stacks for cilia of this strain and obtained the cilium-intensity profiles from the Z-average projection. Additionally, the distal segment of cilia do often overlap, this does create higher intensity. Given our experimental setup, this is unavoidable. Importantly, it does not impact our conclusions.

### Centre of Mass (COM) calculations

The calculation of the COM and the variance around COM was performed in Python using custom scripts. COM was calculated by the formula: 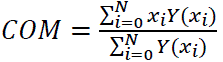

where x is distance, y is the normalized intensity, i is the index of discrete bin’s and N is the total number of bins. To calculate the variance around the COM we used the following formula:

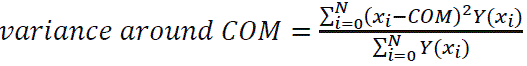

### Kymographs and plots

Kymographs were generated by kymographClear or multiKymographs plug-ins of Fiji/ImageJ^29^. Analysis was performed in MATLAB (The Math Works, Inc., R2021a) with custom written scripts^47^. Plotting was done in Python using custom scripts.

### Single-molecule tracking

Single-molecule tracking was performed as described before^46,47^. In short, movies were acquired at high laser power within a small illuminated region, after photobleaching using the full excitation beam. Movie duration ranged between 5-10 minutes. In case of drift or movement of the worm, images were not analysed further. We estimate the localization precision of individual fluorescent proteins in our imaging conditions to be ∼40 nm (2*σ*), as determined for surface-bound eGFP in our experimental set-up^28^.

### Tracking, spline fitting and defining the ciliary base

Tracking was perform as described previously^26,47^ using a MATLAB-based tracking software, FIESTA (version 1.6.0)^49^. Tracks, corresponding to an event, contain information regarding time (*t_i_*), x and y coordinates (*x_i_*, *y_i_*) and distance moved (*d_i_*), for every frame *i*. Tracks smaller than 12 frames were discarded from further analysis. Erroneous tracks, primarily caused by two (or more) single-molecule events too close to be discriminated, were also excluded from further analysis (apart from spline fitting). Retrograde tracks were observed and used in spline fitting. They are, however, rare because often fluorescence signal that enter the cilia at the base will be bleached before they can become retrograde events. A ciliary coordinate system was defined by interpolating a spline on a segmented line drawn along the long axis of the imaged cilium, visualized via the single-molecule localizations obtained from tracking. A reference point was picked at the base of the characteristic “bone-shaped” structure, at the ciliary base. For all *dyf-5* KO worm lines, this “bone-shaped” structure was not clearly but often there was a cluster of localizations, the proximal part was selected as ciliary base.

All single-molecule localizations were transformed from x– and y-coordinates (*x_i_*, *y_i_*) to ciliary coordinates (*c*_∥_*i*_, *c*_⊥_i_), with *c*_∥_the distance from the reference point along the spline and *c*_⊥_ the distance perpendicular to the spline (*c*_⊥_).

### Classification of directed transport and pausing

Classification was performed as described previously^47^.To obtain a quantitative measure for the directedness of the motion, we used an MSD-based approach to extract the anomalous exponent (α) from *MSD*(*τ*) = 2Γ*τ⍺* (where Γ is the generalized transport coefficient and *τ* is the time lag) along the track, in the direction of motion. α is a measure of the directedness of the motion, α = 2 for purely directed motion, α = 1 for purely diffusive motion and α < 1 for sub-diffusion or pausing. For each data point (*c*_∥_*i*_), we calculate *⍺* in the direction parallel to the spline (*⍺*_∥_*i*_), using a windowed Mean Square Displacement classifier (wMSDc) approach, described in Danné et al^50^. α was calculated analytically, using the following equation: 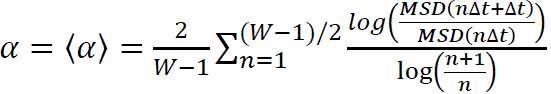; keeping a fixed window (W = 12 frames). Due to the size of the window, all tracks shorter than 12 frames were removed from the analysis. As we here only look at directed motion for calculating velocity, we used a cut of for α ≥ 1.5.

### Velocity measurement

Before calculating the point-to-point velocity, the tracks were smoothened by rolling frame averaging over 10 consecutive time frames, to reduce the contribution due to localization error (typically estimated to be between 10-40 nm, depending on the brightness of the tracked object). The point-to-point velocity at a given localization (*xi*, *yi*) was calculated using the following equation: 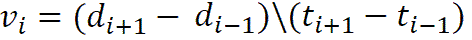

Only velocity corresponding to moving data points (α ≥ 1.5) are displayed in the figures, where we calculated the average velocity and error of the velocity at bins of 100nm along the long axis of the cilia/dendrite, using bootstrapping.

### Estimating average value and error

We used a bootstrapping method to calculate the parameters of a distribution as described before^41^. We randomly selected N measurements from the distribution (with replacement) and calculated the median of the resampled group. We repeated this process 1000 times, creating a bootstrapping distribution of medians. We then calculated the mean (μ) and standard deviation (σ) of this distribution to estimate the parameter and its error. All values and errors are presented as μ ± 3σ.

### Velocity distribution

For histograms showing distribution of velocities over distinct intervals, velocity data was collected over 500 nm intervals along the cilia length and the bin size used is 100nm/s

## Acknowledgements

We thank Dr. Tiago Dantas (University of Porto, Portugal) and his lab, as well current and previous members of our lab for fruitful discussions. We acknowledge financial support from the European Research Council under the European Union’s Horizon 2020 research and innovation programme (Grant agreement no. 788363; “HITSCIL”), Marie Sklodowska-Curie Actions Postdoctoral Fellowship of the European Commission (Project no. 898006; ‘MingleIFT’, A.M.), and from the Netherlands Organisation for Scientific Research (NWO) via an ALW Open Program grant.

## Data availability

The data in this study can be found at https://doi.org/10.34894/BUNQFD

